# Associations between depth and micro-diversity within marine viral communities revealed through metagenomics

**DOI:** 10.1101/551291

**Authors:** FH Coutinho, R Rosselli, F Rodríguez-Valera

## Abstract

Viruses are extremely abundant and diverse biological entities that contribute to the functioning of marine ecosystems. Despite their recognized importance no studies have addressed trends of micro-diversity in marine viral communities across depth gradients. To fill this gap we obtained metagenomes from both the cellular and viral fractions of Mediterranean seawater samples spanning the epipelagic to the bathypelagic zone at 15, 45, 60 and 2000 meters deep. The majority of viral genomic sequences obtained were derived from bacteriophages of the order *Caudovirales*, and putative host assignments suggested that they infect some of the most abundant bacteria in marine ecosystems such as *Pelagibacter*, *Puniceispirillum* and *Prochlorococcus*. We evaluated micro-diversity patterns by measuring the accumulation of synonymous and non-synonymous mutations in viral genes. Our results demonstrated that the degree of micro-diversity differs among genes encoding metabolic, structural, and replication proteins and that the degree of micro-diversity increased with depth. These trends of micro-diversity were linked to the changes in environmental conditions observed throughout the depth gradient, such as energy availability, host densities and proportion of actively replicating viruses. These observations allowed us to generate hypotheses regarding the selective pressures acting upon marine viruses from the epipelagic to the bathypelagic zones.

## Introduction

Viruses are increasingly recognized as important players in the functioning of marine ecosystems[1, 2]. In recent years many efforts were undertaken do describe associations between viral biodiversity and spatial[3], temporal[4], and ecological[5] gradients. The taxonomic composition and functioning of host communities respond to such changes in environmental parameters across such gradients[6, 7]. In response, viruses adapt to those changes to guarantee their survival. The depth gradient of stratified water masses displays marked changes in environmental conditions mainly driven by light availability and temperature[8]. Thus it is an ideal habitat to study associations between environmental parameters, viruses, and their hosts.

In stratified waters, temperature decreases with depth while the concentration of inorganic nutrients increases. The micro-habitat at the thermocline provides photosynthetic microorganisms with ideal conditions of temperature, nutrient availability and light irradiation. The intense proliferation of photosynthetic microbes there leads to a peak of chlorophyll concentration and microbial cell density, known as the deep chlorophyll maximum (DCM). In the stratified water column, the DCM often exhibits the highest densities of prokaryotic cells and viral particles[9, 10]. Moving towards the aphotic zone, the concentrations of inorganic nitrogen and phosphorus increase, but the gradual decrease of light hampers productivity, thus leading to much lower cell densities than observed in the surface or the DCM. Below the DCM both viral and bacterial abundances decrease, and deeper waters of the bathypelagic zone often display the lowest densities of both bacteria and viruses [9, 11].

Previous studies have used metagenomics to assess changes in the taxonomic and functional composition of viral communities throughout depth gradients[4, 12]. Nevertheless, studies addressing patterns of micro-diversity, i.e. accumulation of mutations within genomes, through the stratified water column are lacking. Investigating patterns of micro-diversity can help to elucidate the selective pressures acting upon viral genomes. For example, in co-evolution experiments in which bacteriophages and hosts are cultured together over multiple generations, viruses tend to preferentially accumulate mutations in genes that affect their host range and the productivity of viral particles[13, 14]. These discoveries provided insightful information regarding the processes by which viruses adapt to more efficiently infect their hosts in cultures. Yet no studies have addressed this topic in free-living marine viral communities through culturing-independent approaches. These are necessary because the selective pressures acting on viral genomes in cultures and environmental communities might be drastically distinct.

Here we sought to investigate micro-diversity patterns in the environment to generate hypotheses about the selective pressures acting on marine viral communities throughout the depth gradient. We selected a site at the Mediterranean sea off the coast of Spain during a period of water stratification (October 2015). Seawater samples were retrieved from multiple depths ranging from the epipelagic at 15, 45 (DCM), 60 (DCM) to the bathypelagic at 2000 meters deep, and used for preparing both cellular and viral metagenomes (viromes). Viromes were assembled to obtain complete or partial viral genomes and cellular metagenomes were assembled and binned to obtain metagenome assembled genomes (MAGs). Next, reads from both the viral and cellular fractions were mapped against the assembled viral scaffolds to calculate the level of micro-diversity for each viral protein. Our rationale was that the changes taking place in microbial communities at surface, DCM and aphotic habitats would subject the associated viral communities to different constraints, which would be reflected in the micro-diversity patterns within viral genomes.

## Results

### Assembled viral genomes and predicted hosts

Assembly of viral metagenomes yielded 10,263 genomic sequences of length equal or greater than 5 kbp, within which 133,352 protein encoding genes were identified (Table 1). A total of 7,164 (69.8%) scaffolds were classified as *bona fide* viral sequences based on the annotation of their protein encoding genes (see methods). Among these, 21 scaffolds with length equal or above 10 Kbp (average length = 44 Kbp) and with overlapping ends were identified, which likely represent complete viral genomes. Computational host predictions were obtained for the *bona fide* viral sequences by scanning viral and prokaryote genomes for three signals of virus-host association: homology matches (i.e. long genomic segments sharing high nucleotide identity), shared tRNA genes, and matches between CRISPR spacers and viral sequences. These approaches have previously been benchmarked and shown to provide accurate host predictions, specially at higher taxonomic ranks such as phylum and class[15, 16]. In addition, we manually curated host-predictions by investigating the gene content of viral genomic sequences. Host predictions were obtained for 171 of the *bona fide* viral sequences (Table S1 and Figure 1A). Among those, the majority were predicted to infect *Proteobacteria* (99 sequences), particularly *Alphaproteobacteria* of the genera *Pelagibacter* (52 sequences) and *Puniceispirillum* (38), followed by *Cyanobacteria* (58) of the genera *Prochlorococcus* and *Synechococcus*.

**Table 1:**
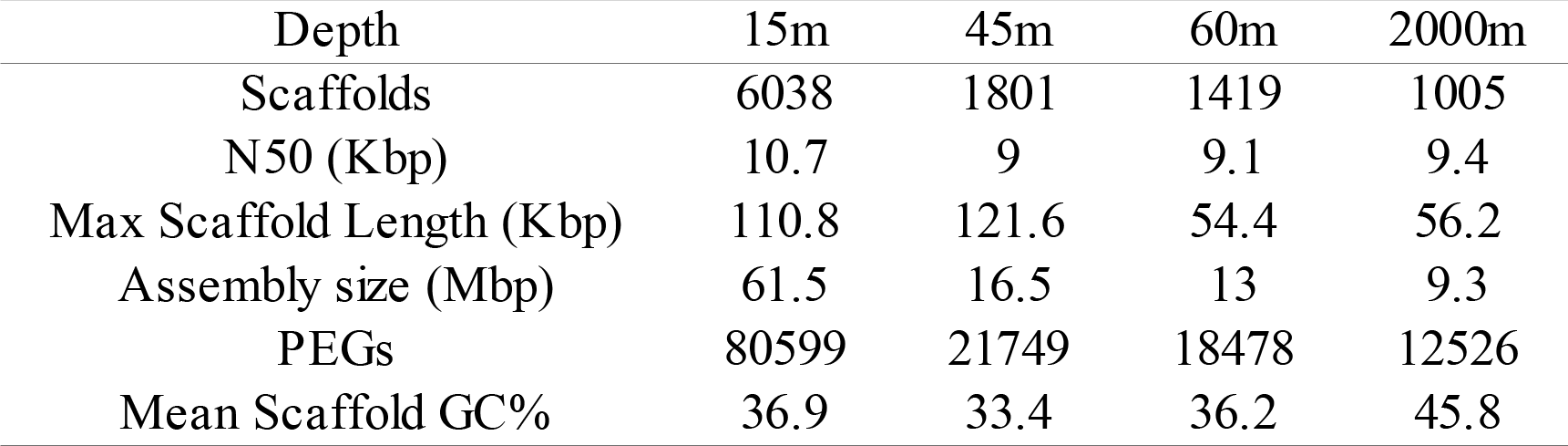
Characteristics of virome assemblies.

**Figure 1:**
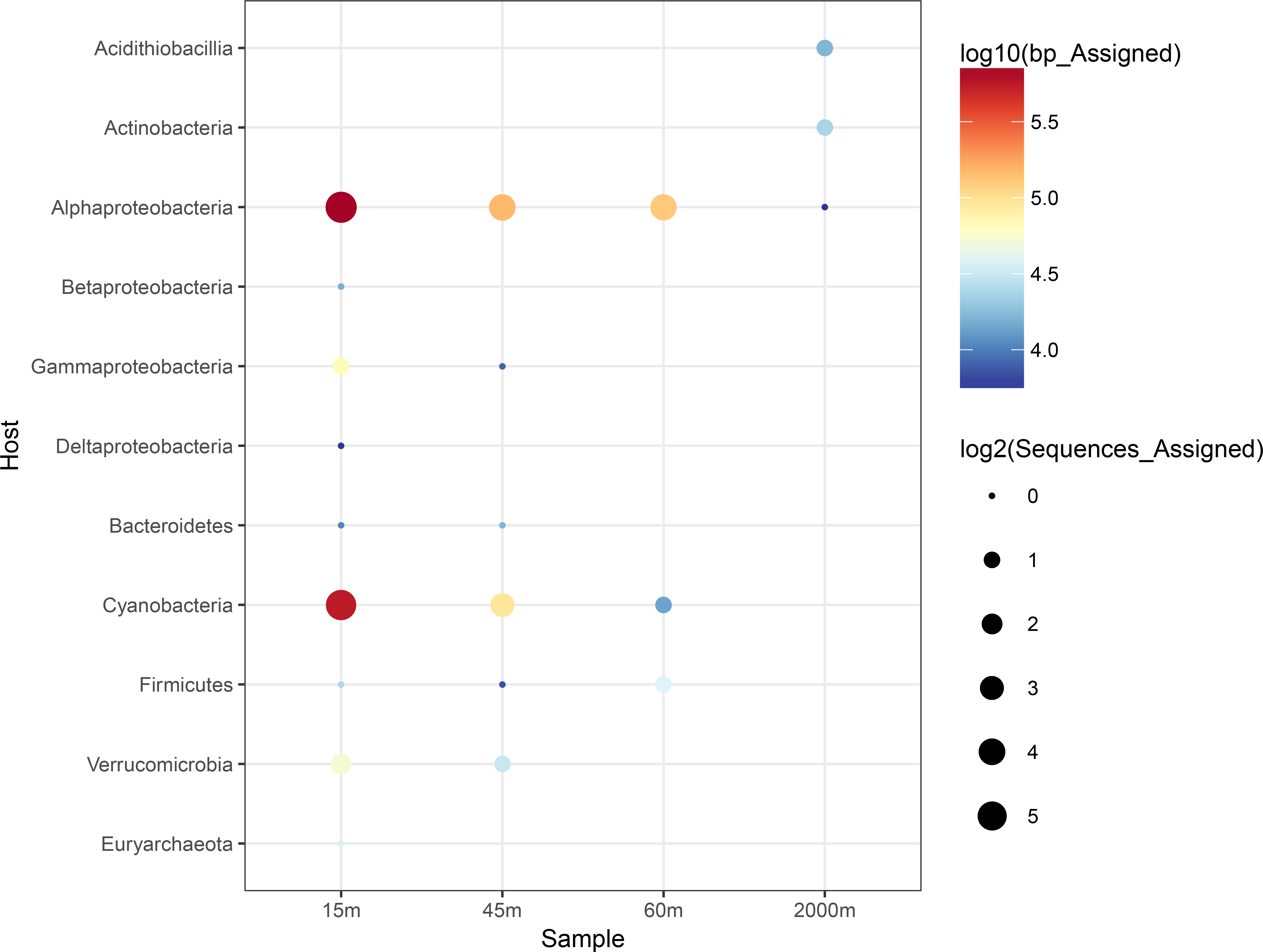
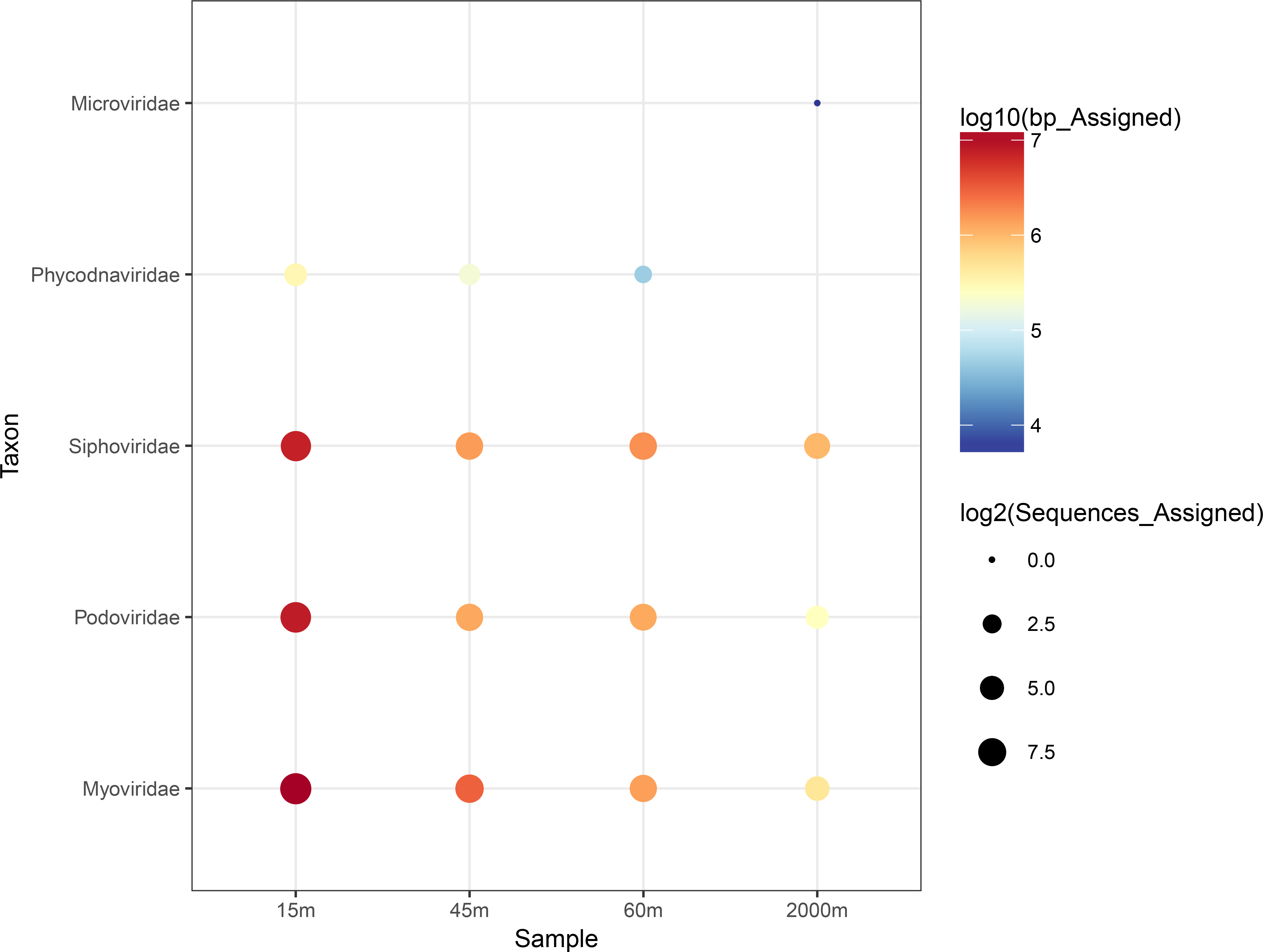
Taxonomic affiliation and predicted hosts of at the *bona fide* viral scaffolds. A) Bubble plot depicting computational host predictions obtained for viral scaffolds. B) Bubble plot depicting taxonomic assignments of the scaffolds based on percentage of matched proteins and average amino acid identity to protein sequences from viral families in the NCBi-nr database.

Taxonomic classification of the assembled scaffolds identified most of them as tailed bacteriophages from the order *Caudovirales* (Figure 1B), specifically as members of the families *Myoviridae*, *Podoviridae* and *Siphoviridae*. Some of the scaffolds from the epipelagic samples were classified as *Phycodnaviridae*, viruses that infect Eukaryotic algae. Scaffolds annotated as *Microviridae* bacteriophages were exclusively retrieved form the bathypelagic sample.

### Viral community composition

Grouping viral abundances according to predicted host revealed differences among samples of the depth gradient (Figure 2A). Scaffolds predicted to infect *Proteobacteria* were among the most abundant in all depths with abundances ranging from 0.5% to 2.4% of mapped reads. Scaffolds predicted to infect *Cyanobacteria* and *Euryarchaeota* displayed their highest abundances at the15m and 45m samples while those predicted to infect *Bacteroidetes* were abundant only at the 45m sample. The 2000m displayed a unique profile with abundant scaffolds predicted to infect *Firmicutes* and *Actinobacteria*.

**Figure 2:**
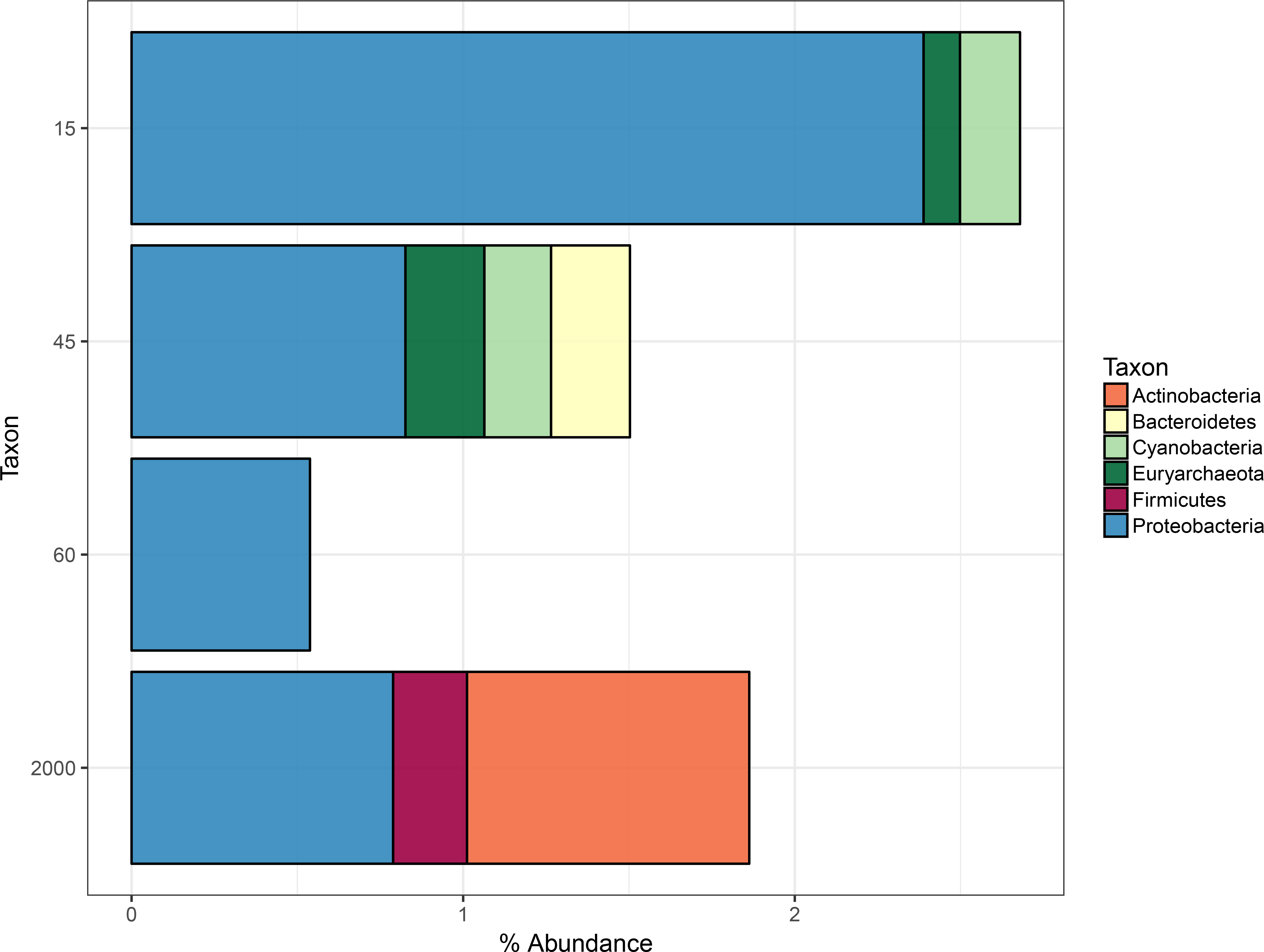
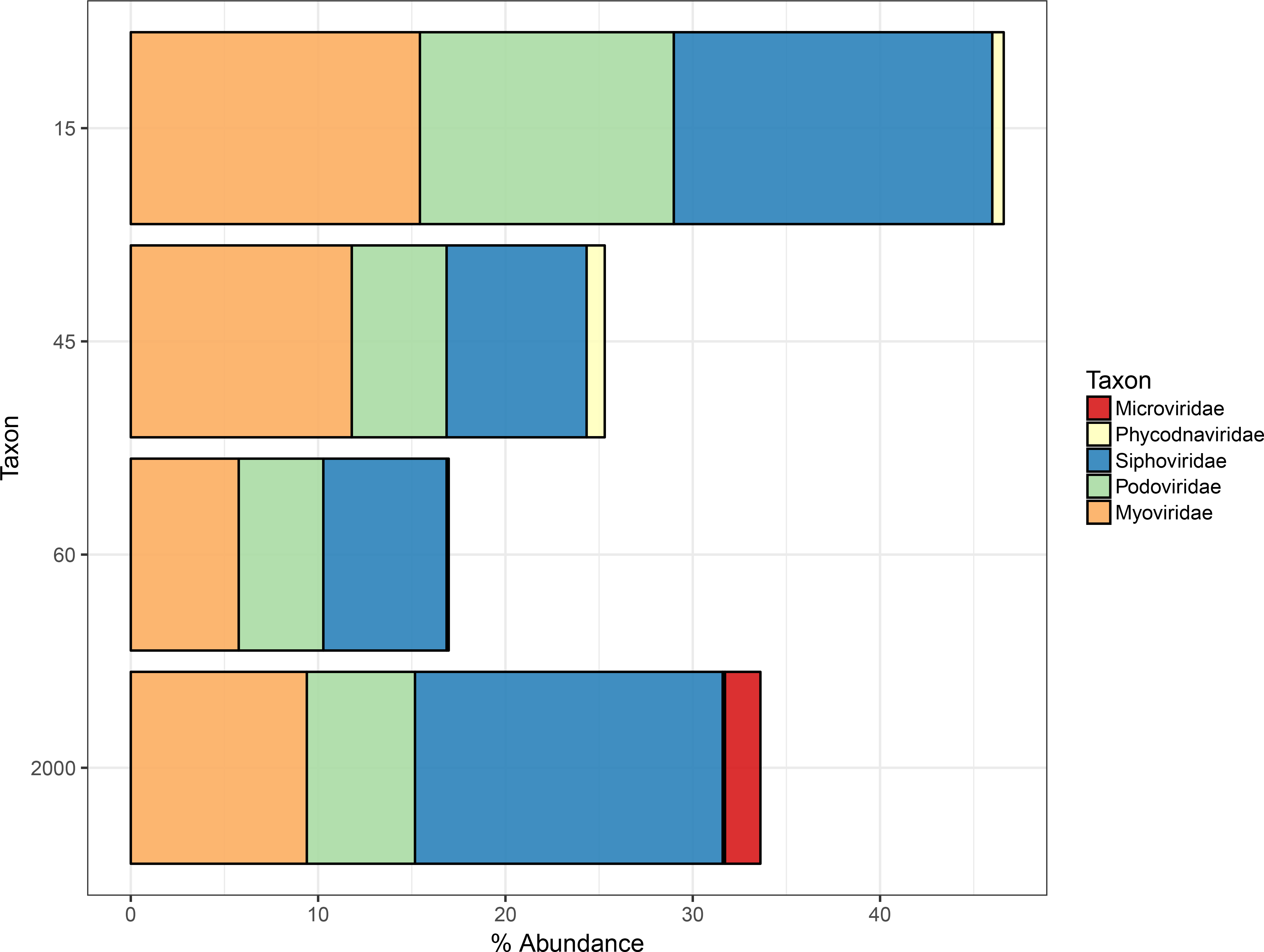
Viral community composition profile across the depth gradient. Bar plots depicting abundances in viromes based on raw read annotation against the database of assembled viral scaffolds. A) Scaffold abundances were grouped according to the phylum level putative hosts of viral scaffolds. B) Scaffold abundances were grouped according to the family level taxonomic affiliation of viral scaffolds. Only taxa that displayed relative abundances equal or above 0.1% are shown.

Previous investigation of the metagenomes from the cellular fraction revealed shifts in taxonomic composition of prokaryotic communities throughout the depth gradient[8]. These were dominated, at all depths, by *Proteobacteria*, mostly from the classes *Alphaproteobacteria* and *Gammaproteobacteria*. The taxonomic composition of viral communities also displayed shifts according to depth (Figure 2B). The families of tailed bacteriophages *Myoviridae*, *Podoviridae* and *Siphoviridae* within the order *Caudovirales* were dominant in all samples, and together accounted for 15% to 45% of the annotated reads. Bacteriophages from the family *Microviridae* were abundant in the bathypelagic sample only, while eukaryotic viruses from the family *Phycodnaviridae* were detectable only at the epipelagic samples, although at lower abundances.

### Mediterranean viruses actively replicating in the cellular fraction

Read mapping revealed that many of the viral scaffolds assembled from viromes could also be detected in the cellular metagenomes (Figure 3A). We assumed that the viral sequences that were abundant in the cellular metagenomes are derived from actively replicating viruses undertaking lytic infections, which lead to high copy numbers of their genomes inside host cells. Alternatively, viral sequences in the cellular fraction could be the result of lysogenic infections. Yet those are not expected to produce the high copy numbers of viral genomes inside host cells that could lead to the observed abundance patterns.

**Figure 3:**
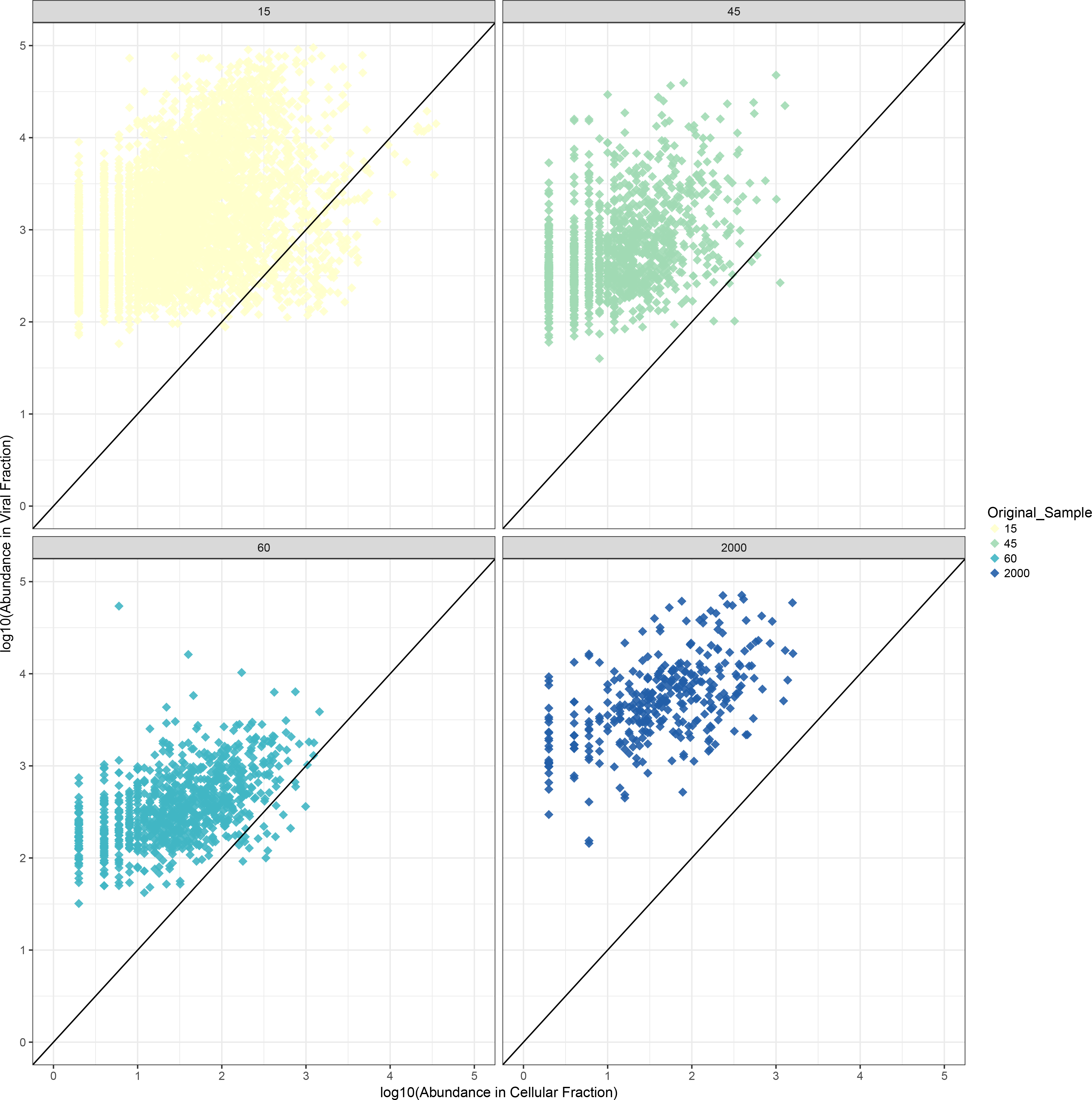
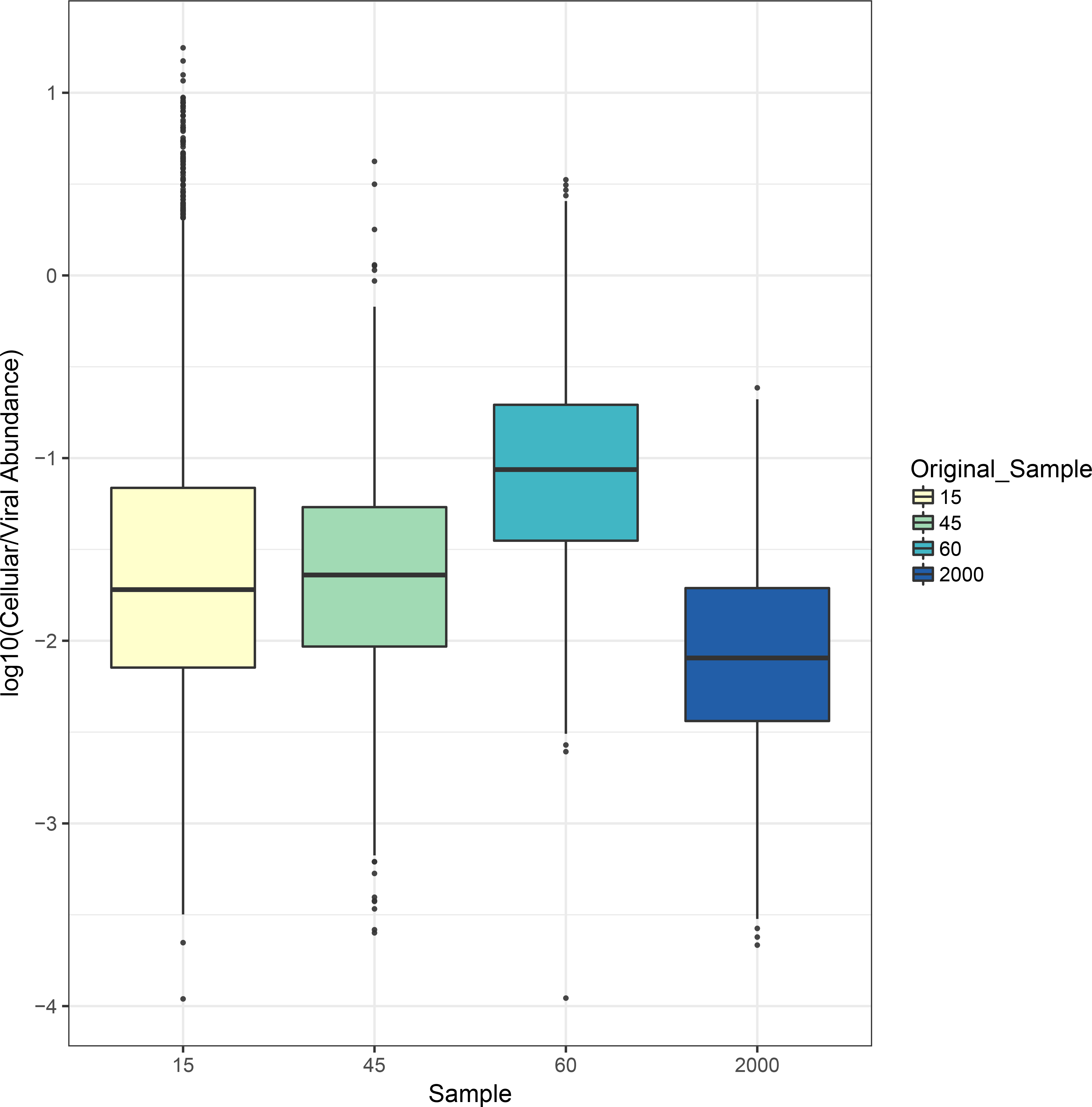
Viral scaffold abundances in viral and cellular metagenomes from the depth gradient. A) Scatter-plots depicting the relative abundances of viral sequences in the viral (Y axis) and cellular (X axis) metagenomes. B) Boxplots depicting the ratio between abundances in the cellular and viral fractions for each sample. Boxes depict the median, the first and third quartiles. Whiskers extend to 1.5 of the interquartile ranges. Outliers are represented as dots above or below whiskers.

Abundances of viral sequences in the cellular fraction differed between samples. The average ratios of cellular/viral abundances were highest for the 45 and 60m samples, followed by 15m and lastly the 2000m sample (Figure 3B). Likewise, the abundance of raw reads annotated as viral in the cellular fraction metagenomes followed the same trend. Thus, there were more viruses actively replicating at the DCM samples than at any other depth, followed by the 15m sample and lastly the 2000m sample, which displayed the lowest proportion of actively replicating viruses. In addition, the DCM samples displayed the lowest values for the Shannon diversity index (5.55 and 5.61), while these values were higher for the 15m (7.21) and 2000m (7.26) samples. The high proportion of actively replicating viruses, and the low Shannon diversity observed at the DCM suggest that the intense viral replication taking place at these depths lead to a highly clonal community, with many nearly-identical viral genomes co-existing at high densities.

### Levels of micro-diversity shift throughout the depth gradient and across functional categories

We evaluated micro-diversity patterns by measuring the pN/pS ratios of protein encoding genes identified in the *bona fide* viral scaffolds. The pN/pS ratio is a measure analogous to dN/dS that does not require specific haplotypes to be identified, and therefore can be applied to metagenomic datasets to provide a population level measure of micro-diversity[17–19]. Briefly, reads from the metagenomes were mapped to the assembled scaffolds to detected mutations, specifically single nucleotide polymorphisms. Next, pN and pS were calculated by respectively dividing the observed counts of non-synonymous and synonymous mutations by the expected frequencies of these mutations under a neutral model.

The majority of proteins displayed pN/pS values below 1, regardless of sample, meaning that the frequencies of non-synonymous mutations was below that which was expected by chance. Thus purifying selection was a major driving force regulating frequencies on mutations among viral genes. Nevertheless, 117 proteins displayed pN/pS above 1 in the cellular fraction metagenomes, and 1,092 in the viral fraction metagenomes. Most of these proteins were retrieved from the 15m sample (755), followed by 2000m (239), 45m (148) and 60m (67) samples. Although the majority of these genes had no assigned functions, some were identified as: recombinase/nuclease proteins (21), oxygenases (17), lysins (16), methylases (13), and tail fibers (11).

We observed a negative association between depth and the median pN/pS ratio of each sample (Figure 4A). The highest median of pN/pS values was observed for the 2000m sample, followed by 60m, 45m and lastly the 15m sample. These trends of pN/pS and depth were observed for viral sequences detected in the metagenomes from both the viral and cellular fractions. Because the coverage of proteins in the viral fraction metagenomes was much higher and spanned many more of the viral proteins, we focused subsequent analysis of pN/pS using viral fraction metagenomes only.

**Figure 4:**
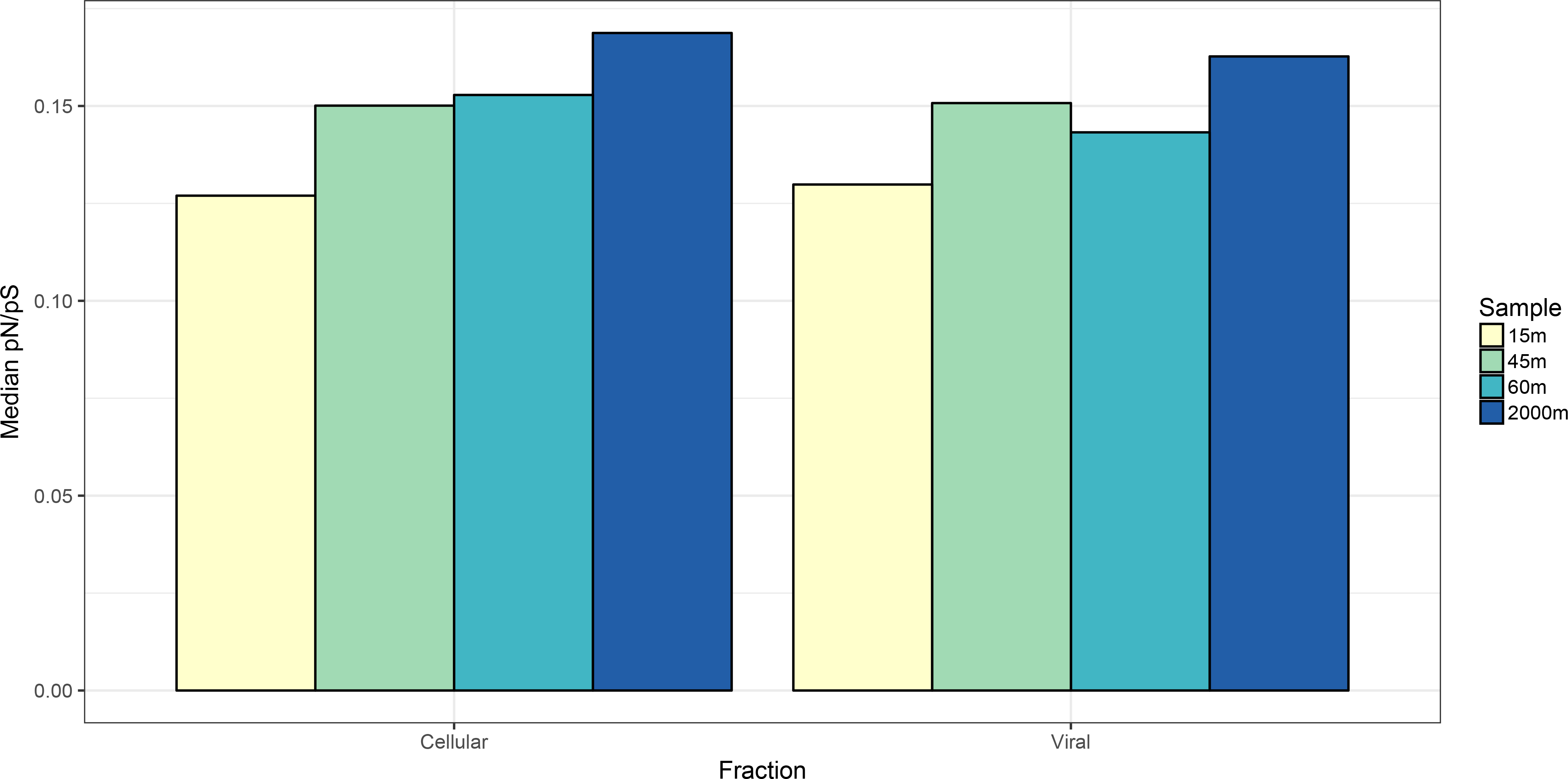
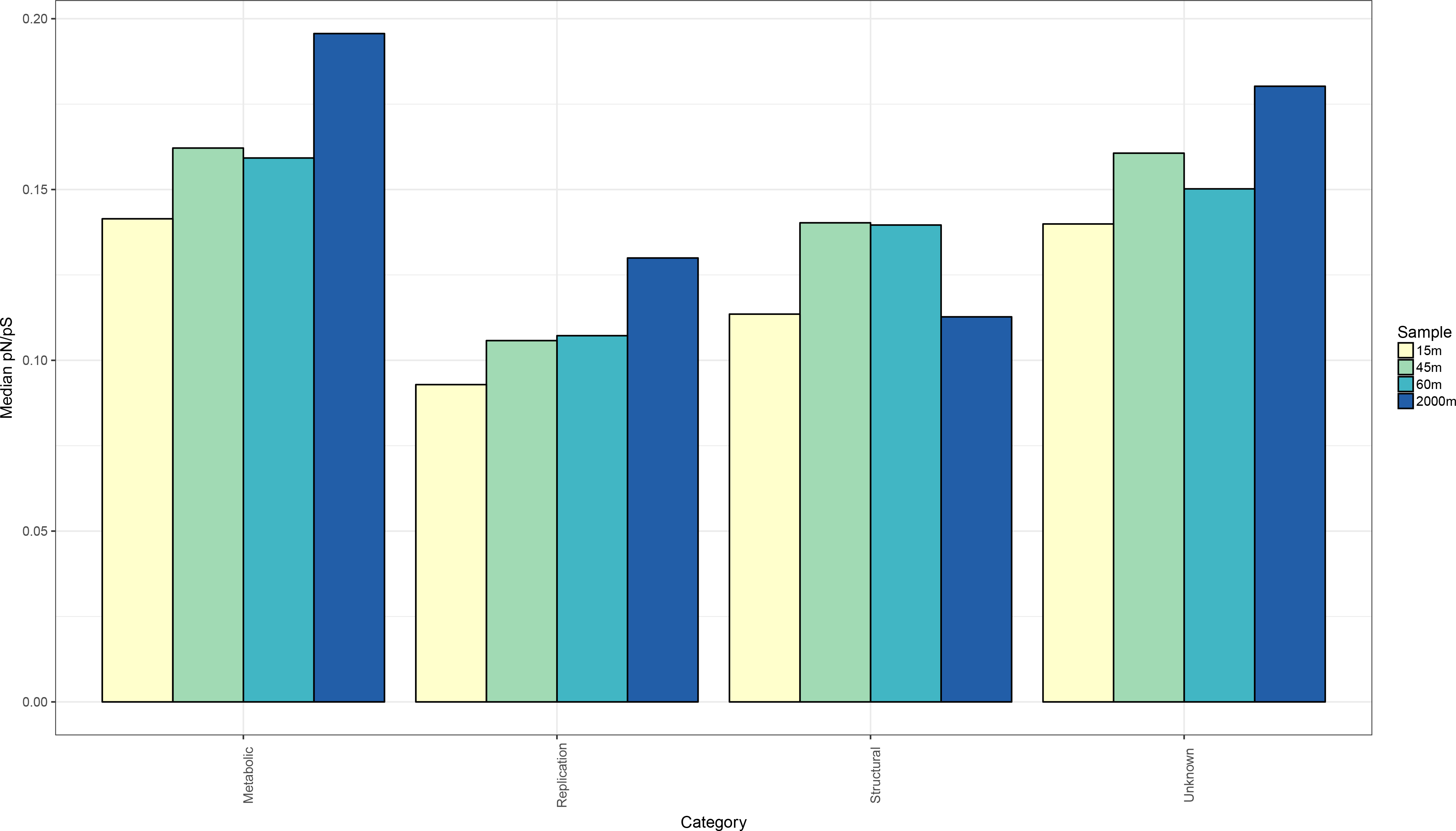
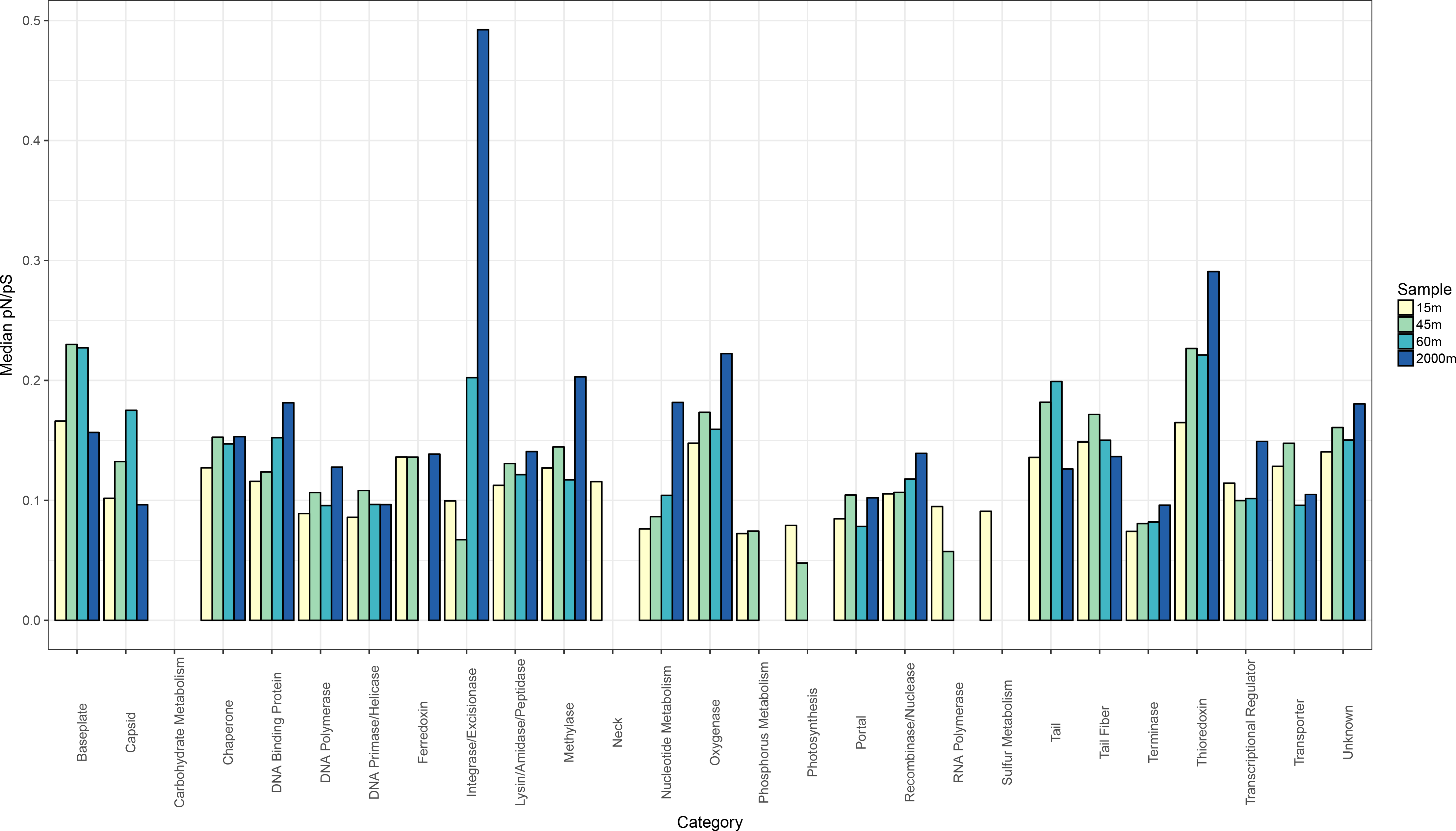
pN/pS values of viral genes differ among functional categories.. A) Barplots depict the median pN/pS values of the functional categories of each sample for the cellular and viral fractions B) Median pN/pS values of proteins grouped by sampling site and broad functional category for the viral fraction only. C) Median pN/pS values of proteins grouped by sampling site and specific functional category for the viral fraction only. Only proteins derived from the set of *bona fide* viral sequences were included in these analyses. When calculating medians only proteins that displayed pN and pS values above 0 were included. Also, only proteins with a total number of polymorphic sites equal or above 1 and percentage of polymorphic sites equal or above 1% were included, so to avoid estimating pN/pS values based only on a small fraction of protein length. Median values obtained from less than three proteins were omitted.

Due to the many unknown proteins present in marine viral genomes, our capacity to annotate these genes and predict their function is limited[20]. Nevertheless, we observed marked differences of median pN/pS ratios among proteins according to functional categories (Figures 4B and 4C). Genes involved in genome replication (e.g. DNA polymerase, DNA primase and genes of the nucleotide metabolism) displayed the lowest median pN/pS values compared to other categories. Structural viral proteins (e.g. capsid, neck and tail) showed intermediate median pN/pS values. Finally, proteins associated with altering host metabolism (e.g. ferrochelatases, thioredoxins and oxygenases) displayed the highest median pN/pS values. A positive association between pN/pS and depth was also observed when grouping proteins according to broad functional categories (Figure 4B). A notable exception was the median pN/pS ratio of structural proteins, which was highest for the DCM samples.

These differences of pN/pS among functional categories are associated with their roles during the viral infection cycle. Genes involved in genome replication must operate at high fidelity and efficiency, thus deleterious non-synonymous mutations in these proteins are readily removed from the population by purifying selection. Meanwhile, structural proteins are fundamental for adequate particle assembly, encapsulation of the viral genome, and host recognition. Deleterious mutations in structural genes can also compromise viral infections, but not as much as errors during genome replication. Finally, metabolic genes are responsible for re-directing host metabolism towards pathways that favour viral particle production[21, 22]. Thus, lower efficiency of metabolic genes due to deleterious mutations is likely to reduce viral productivity but not to compromise it as much as deleterious mutations in the genome replication or structural modules.

### The DCM is a micro-diversity hot-spot for viral receptor binding proteins

The DCM samples displayed the highest median pN/pS values for structural proteins (Figure 4B). Specifically, structural proteins that encoded baseplate, capsid, tail, and tail fiber genes displayed pN/pS values higher than their counterparts in the remaining samples (Figure 4C). Interestingly all of these proteins either are or interact directly with receptor binding proteins that mediate host recognition, a fundamental step for successful viral infection[23, 24]. The enhanced pN/pS observed for these genes at the DCM provides evidence that this habitat is a micro-diversity hot-spot for viral receptor binding proteins.

Adaptation to sub-optimal hosts is a major driver of genomic diversification for viruses, which is associated with the quick accumulation of non-synonymous mutations in tail fiber proteins[14]. A single nucleotide polymorphism in tail fiber gene can be sufficient to alter viral host range[13, 25]. Consistent with those findings, we observed multiple cases of tail proteins in which non-synonymous mutations were concentrated in small segments of these gene (Figure 5). These sites that accumulate non-synonymous mutations at higher frequencies than the other codons are likely those that confer a selective advantage to the virus at their specific habitat according to the availability of hosts. These trends are consistent with a scenario where, on the one hand, positive selection acts on tail fiber proteins to expand host range, while on the other hand, purifying selection removes mutations from other sites where they cause loss of function or restrict the host range instead of expanding it[14, 26].

**Figure 5:**
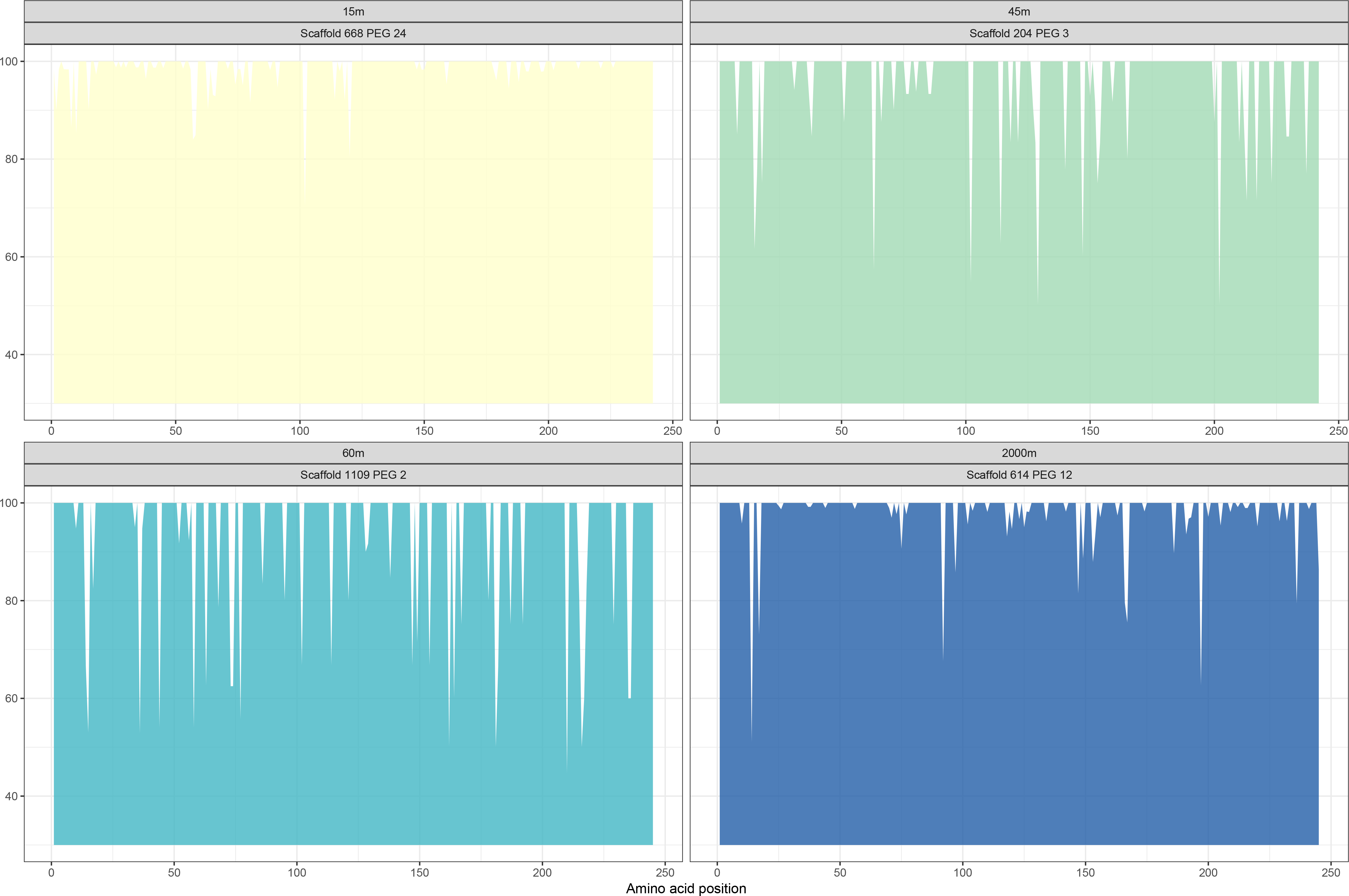
Micro-diversity patterns within a group of homologous tail proteins. X axis depicts the amino acid position along proteins. Y axis depicts the frequency of the reference amino acid among the viral population from each sample. Valleys in the plot represent areas that concentrate non-synonymous mutations, possibly driven by positive selection favouring mutations that modify or expand host-range.

## Discussion

### Different selective pressures determine levels of micro-diversity throughout the depth gradient

Major changes take place among prokaryotic and viral communities throughout the depth gradient, affecting their taxonomic composition and virus-host interactions [4, 8, 9, 27, 28]. These differences in cell densities and frequency of replication events impact micro-diversity because the rate at which viral genomes accumulate mutations is density dependent, meaning that they adapt faster in conditions with higher host density, in which more infection events take place[30]. Our results demonstrated that the DCM viral communities had the highest proportions of actively replicating viruses but were the least diverse. We propose that this scenario leads to intense intra-species competition between viruses for suitable hosts, creating a selective pressure that favours viruses with mutations in receptor binding proteins which provide them with a different host-range, allowing them to exploit a distinct niche (Figure 6). The high micro-diversity observed among receptor binding proteins and the clonal populations observed within DCM samples suggests that many strains of viruses with distinct host ranges co-exist at this habitat. It follows that host strains with different patterns of viral-susceptibility are also co-existing in these sites. This is in agreement with the constant diversity theory[29], which postulates that the trade-offs between ecological fitness and viral susceptibility are responsible for avoiding that a single bacterial clone dominates the community through clonal sweeps, thus preserving the taxonomic and functional diversity of these communities[13, 26].

**Figure 6:**
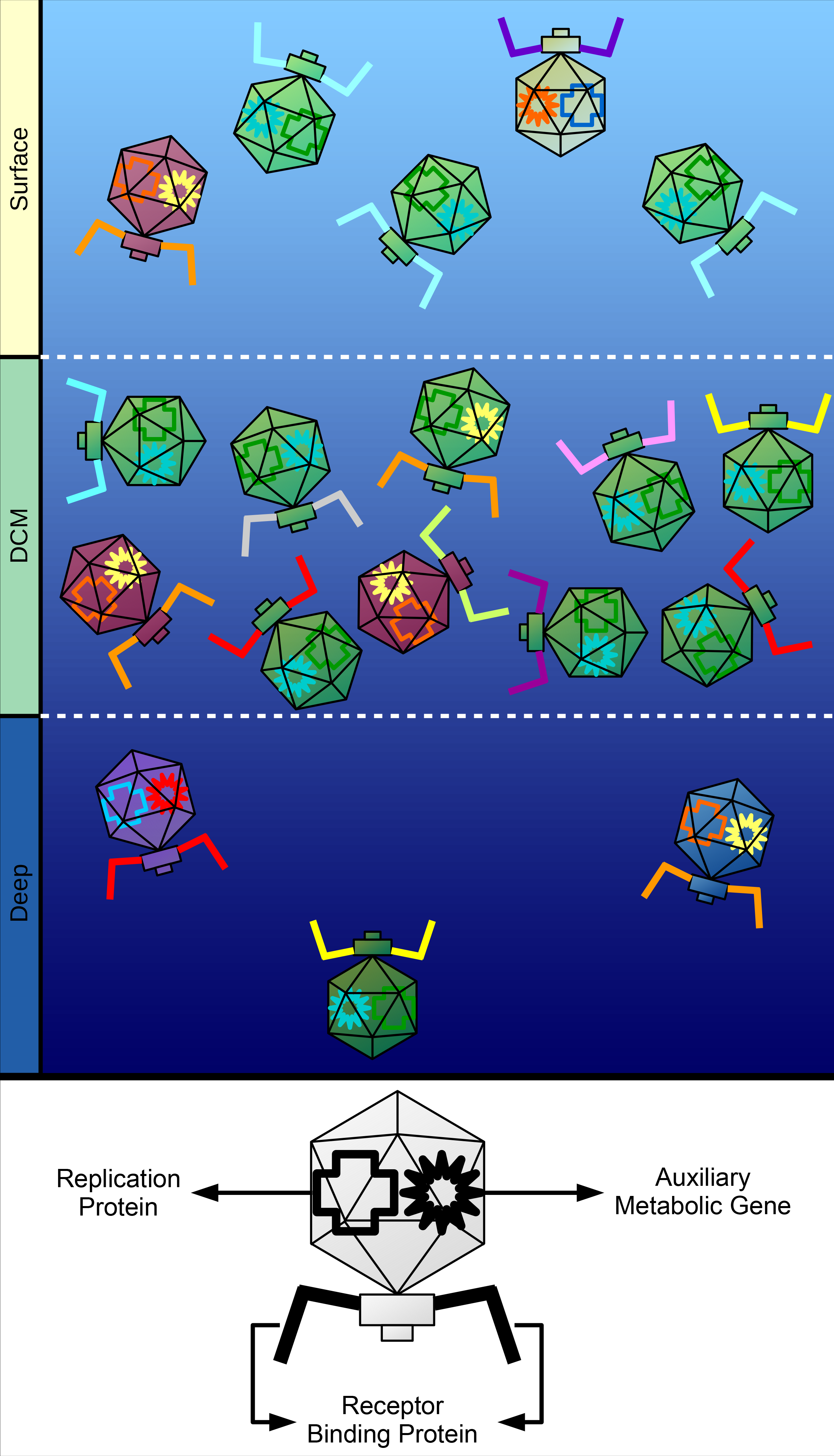
Conceptual model summarizing the observed patterns of micro-diversity in marine viral genomes across the depth gradient. Different capsid colours represent different viral species. Different colours for receptor binding proteins, auxiliary metabolic genes and replication proteins represent different isoforms of the same protein created by non-synonymous mutations. Surface samples have intermediate densities of viral particles and intermediate species diversity, this sample displayed the lowest degree of micro-diversity for all functional categories. DCM samples have the highest density of viral particles but the lowest species diversity. These samples displayed the highest degree of micro-diversity among receptor binding proteins. Deep samples have the lowest density of viral particles but highest species diversity. This sample displayed the highest degree of micro-diversity among metabolic and replication proteins.

Meanwhile, a different pattern was observed for the bathypelagic sample. In this habitat both viral and cell densities are much lower than in the DCM or the surface[9, 11]. Due to the lower availability of hosts at 2000 meters, chance encounters between viruses and hosts are expected to occur less often. Thus, less infection events take place at 2000m compared to shallower depths with higher host densities, as evidenced by the differences in abundances of viruses actively replicating in the cellular fraction. Interestingly, the bathypelagic sample displayed the highest Shannon diversity but lowest proportion of actively replicating viruses. This finding suggests that in the energy-limited bathypelagic zone, intra-species competition for hosts is expected to be less relevant than it is at the DCM, where a highly clonal population with high density was observed. Instead, the major constraint faced by viruses at this depth could be the efficient production of viral progeny, since in this scenario a lower reproductive fitness is more likely to lead to local extinction than in the highly productive conditions of the euphotic zone. Consistent with that, we observed the highest pN/pS values of proteins encoding metabolic functions (e.g. oxygenases and thioredoxin) and transcriptional regulators in the 2000m sample. We postulate that the higher micro-diversity observed among these genes in the bathypelagic sample is evidence of positive selection acting on proteins that increase the capacity of viruses to generate progeny by using a diverse array of auxiliary metabolic genes and transcriptional regulators to fine-tune host metabolism to enhance the production of viral particles under conditions of low energy availability and productivity (Figure 6).

### Micro-diversity patterns differ between pure cultures and environmental samples

In laboratory experiments of phage-bacteria co-evolution, mutations usually accumulate in genes involved in host specificity such as tail proteins[14, 31]. In contrast, we observed a broader distribution of mutations that spanned all functional categories within viral genomes. We attribute this to the differences between the selective pressures imposed over viruses in co-evolution experiments versus in the environment. In the former, the only selective pressure is to effectively infect one single host derived from a clonal population. In the latter, viruses have a multitude of hosts available, each with their specific viral receptors and resistance mechanisms (e.g. CRISPR and restriction modification systems). In cultures, once resistance mutations appear their prevalence quickly rises within bacterial populations[14]. In the environment, the frequency of resistance mutations is simultaneously regulated by a trade-off of viral resistance and the fitness cost brought by these resistance mutations[26]. These differences in selective pressures faced by viruses in environmental communities is likely to lead to the accumulation of mutations throughout the entirety of viral genomes, and not just at the sites associated with host recognition and infection.

### Concluding remarks

Light, depth and temperature are main factors structuring the taxonomic and functional composition of marine viral communities[5, 32]. These variables are major determinants of the energy available across the ecosystem, and they shift drastically throughout the depth gradient from which our samples were retrieved[8]. Our data shows that these parameters not only shape the taxonomic composition of viral communities but also influence how the genomes of these viruses accumulate mutations and evolve. To our knowledge this is the first study assessing patterns of micro-diversity within marine viromes. The obtained results allowed us to postulate hypotheses about the selective pressures acting upon marine viruses from the community to the amino acid level. Furthermore, we demonstrated that the frequencies of non-synonymous mutations differed among functional categories and depth. Finally, free-living viruses displayed patterns of mutation accumulation different from those observed in laboratory conditions, which has important implications for how the latter should be interpreted. Here we set a stepping stone for investigating patterns of micro-diversity among environmental viral communities. Further research will be necessary to determine if the patterns presented here are also present in other marine habitats as well as different ecosystems (such as host-associated, freshwater and soils), and to determine the driving forces behind them.

## Materials and Methods

### Sampling and sample processing

Four samples from different depths, 15, 45, 60 and 2000 meters were collected on October 15^th^ 2015 from aboard the research vessel “Garcia del Cid” [8]. The sampling site was located at approximately 60 nautical miles off the coast of Alicante, Spain, at 37.35361° N - 0.286194° W. Sea water samples were filtered for Eukaryote and Prokaryote fractions through 20 μm, 5 μm and 0.22 μm pore size polycarbonate filters (Millipore). Two technical replicates (50 L for each depth) were ultra-filtered on board through a Millipore Prep/Scale-TFF-6 filter, yielding 250 ml of viral concentrate stocks. Each stock was purified through Sterivex 0.22 filters (Millipore), stored at 4°C and subsequently reduced to 1,5 mL using Ultra-15 Centrifugal Filter Units (Amicon).

To minimize the carry-over of free-residual nucleic acids, stocks were treated with 2,5 U of DNase-I at 37°C for 1h, followed by inactivation with EDTA (0,5 mM). Total viral DNA was extracted with PowerViral Environmental RNA/DNA Isolation Kit (MoBio). Quality and quantity of extracted DNA were determined using the ND-1000 Spectrophotometer (NanoDrop, Wilmington, USA) and Qubit Fluorometer (Thermofisher). The absence of prokaryotic DNA was tested through PCR using 16S universal primers on aliquots from each sample. Multiple Displacement Amplification (MDA) was performed using Illustra GenomiPhi V2 DNA Amplification Kit (GE Healthcare, Life Sciences).

### Sequencing, Assembly and Binning

Metaviromes were sequenced using Illumina Hiseq-4000 (150 bp, paired-end reads) by Macrogen (Republic of Korea). Reads from metaviromes were pre-processed using Trimmomatic[33] in order to remove low-quality bases (Phred-quality score of 20 in 4-base sliding windows) and reads shorter than 30 bases. Each metagenome was individually assembled through SPAdes[34] using default parameters for the metagenomic mode. Sequences shorter than 5 Kbp were discarded. Both raw reads and assembled scaffolds were deposited at ENA under project ERP113162. Taxonomic and functional annotation of proteins were performed by querying PEGs against the NCBI-nr database using Diamond[35], and against the pVOGs[36] database using hmmer[37].

Scaffolds from the cellular fraction of the 2000m sample were binned with MetaBat[38] to obtain Metagenome Assembled Genomes (MAGs) of Bacteria and Archaea. Quality of MAGs was assessed through CheckM[39]. MAGs were manually curated to improve completeness and reduce potential contamination. Protein encoding genes were identified using the metagenomic mode of Prodigal[40].

### Computational host prediction

Putative hosts were assigned to viral scaffolds through homology matches, CRISPR spacers and shared tRNAs as previously described[41]. These were performed using two datasets: The NCBI RefSeq genomes of Bacteria and Archaea (June 2017 release), and the MAGs obtained from the binning of scaffolds from the cellular fraction metagenomes obtained from the same samples from which the viromes were derived[8]. Putative hosts were manually assigned for sequences that displayed high similarity to RefSeq bacteriophage genomes as measured by proportion of shared genes and synteny between genomes. Ambiguous host predictions, i.e., derived from viral sequences predicted to infect more than a single taxa were removed from further analyses.

### Abundance profiles and Micro-diversity analysis

Sequencing reads from the cellular and viral metagenomes were mapped to assembled viral scaffolds using Bowtie2 in sensitive-local mode[42]. The number of reads mapped was used to estimate the relative abundances of the viral sequences in both fractions. To estimate mutational frequencies on viral genomes, raw reads were mapped to assembled scaffolds using the sensitive-mode of Bowtie2. Next, the generated bam files were analysed through Diversitools (http://josephhughes.github.io/DiversiTools/) to obtain counts of synonymous and non-synonymous mutations in each protein. Codon mutations were only considered valid if they were detected at least 4 times, in at least 1% of the mapped reads, and if the codon coverage was equal or above 5x. Only the mutations that passed the aforementioned filters were considered to estimate the percentage pN/pS ratios, which were calculated as described in [19].

## Acknowledgments

FHC, RR and FRV were supported by the grants “VIREVO” CGL2016-76273-P [AEI/FEDER, EU], (co-funded with FEDER funds); Acciones de dinamización “REDES DE EXCELENCIA” CONSOLIDER-CGL2015-71523-REDC from the Spanish Ministério de Economía, Industria y Competitividad. FHC was supported by APOSTD/2018/186 Post-doctoral fellowship from Generalitat Valenciana.

